# Effects of BMP-2 dose and delivery of microvascular fragments on healing of bone defects with concomitant volumetric muscle loss

**DOI:** 10.1101/428359

**Authors:** Marissa A. Ruehle, Laxminarayanan Krishnan, Casey E. Vantucci, Yuyan Wang, Hazel Y. Stevens, Krishnendu Roy, Robert E. Guldberg, Nick J. Willett

## Abstract

Traumatic composite bone-muscle injuries, such as open fractures, often require multiple surgical interventions and still typically result in long-term disability. Clinically, a critical indicator of composite injury severity is vascular integrity; vascular damage alone is sufficient to assign an open fracture to the most severe category. Challenging bone injuries are often treated with bone morphogenetic protein 2 (BMP-2), an osteoinductive growth factor, delivered on collagen sponge. Previous studies in a composite defect model found that a minimally bridging dose in the segmental defect model was unable to overcome concomitant muscle damage, but the effect of BMP dose on composite injuries has not yet been studied. Here, we test the hypotheses that BMP-2-mediated functional regeneration of composite extremity injuries is dose dependent and can be further enhanced via co-delivery of adipose-derived microvascular fragments (MVF), which have been previously shown to increase tissue vascular volume. Although MVF treatment did not improve healing outcomes, we observed a significant BMP-2 dose-dependent increase in regenerated bone volume and biomechanical properties. While high dose BMP-2 delivery can induce heterotopic ossification (HO) and increased inflammation, the maximum 10 μg dose used in this study did not result in HO and was associated with a lower circulating inflammatory cytokine profile than the low dose (2.5 μg) group. These data support the potential benefits of an increased, though still moderate, BMP-2 dose for treatment of bone defects with concomitant muscle damage. Future work to improve vascularization may further improve healing.

## INTRODUCTION

Composite tissue injuries present a compelling clinical challenge to both military and civilian trauma populations. Of the six million fractures that occur in the U. S. each year, approximately 10% are open fractures, which are characterized by concomitant soft tissue damage. These injuries experience a complication rate about twice that of closed fractures, with complications including infection, malunion, and non-union ^1^. As a result, open fractures are often subject to multiple treatment strategies, repeated surgeries, and high direct and indirect costs ^2^. Despite these multiple interventions, nearly two-thirds of patients remain significantly disabled long term (at seven years post-injury, and likely beyond) ^3^.

One treatment strategy often employed in challenging bone healing scenarios is the delivery of bone morphogenetic protein-2 (BMP-2) on a collagen sponge. While most commonly used in cases of spinal fusion ^4; 5^, BMP-2 has demonstrated clinical success in long bone fracture cases for over 15 years ^6^. However, BMP-2 has also been linked to side effects such heterotopic bone formation and inflammation ^7^, both of which can be exacerbated by orthopaedic trauma ^8^. Segmental bone defect healing is known to exhibit a dose response to BMP-2 ^9^; however, a minimally bridging dose for a segmental bone defect was unable to consistently bridge same-sized bone defects in a composite bone and muscle defect model ^10^. Thus, the effect of BMP-2 dose is of particular interest in traumatic composite defects.

The Gustilo classification system scores open fractures from class I-IIIC in terms of increasing severity. Assignment to broad classes I-III is determined by increasing size of soft tissue loss, but the hallmark of the most severe category IIIC is vascular damage – regardless of the degree of soft tissue damage ^11^. The revascularization process is a rate-limiting step of the wound healing^12^ and plays a crucial role in the clinical success of treatments for open fractures. Further, it is well established that angiogenesis and osteogenesis are linked ^13^ and that impaired revascularization impairs fracture healing in pre-clinical animal models ^14^. Numerous therapeutic strategies targeting revascularization of bone have been employed, largely pre-clinically, including growth factor and cell delivery ^15–19^.

One promising approach is the use of microvascular fragments (MVF), multicellular segments of mature vasculature that form angiogenic networks in vitro ^20^. Due to their pre-patterned tubular structure and retention of vascular support cells, MVF are attractive candidates for tissue engineering applications^21^, MVF have been shown to inosculate with host vasculature^22;23^ increase tissue vascular volume^23;24^ and improve tissue grafting in pre-clinical models^25;26^. MVF are most commonly obtained from adipose tissue and therefore have potential clinical utility in an autologous application.

We have previously developed a pre-clinical small animal model of composite bone-muscle injury consisting of a critical size femoral defect with an overlying volumetric muscle loss ^10^. Our model recapitulates the clinical observation of impaired bone healing when concomitant soft tissue injury is present, and this has also been seen in similar pre-clinical models of composite tissue injury^10; 27–29^. In this study, we tested the hypotheses that BMP-2-mediated functional regeneration of composite extremity injuries is dose dependent and can be further enhanced via co-delivery of adipose-derived microvascular fragments (MVF), which have been previously shown to increase tissue vascular volume.

## METHODS

### MVF Isolation and Construct Preparation

MVF were isolated as previously described ^20^. Briefly, epididymal fat pads were harvested from Lewis rats, minced, and digested with a collagenase solution for 7 minutes at 37 °C. MVF were obtained through selective filtration to retain tissue between 50–500 μm. To prepare surgical constructs, MVF were suspended at a density of 80,000 MVF/mL in αMEM (ThermoFisher Scientific; Waltham, MA) along with BMP-2 (Pfizer, Inc.; New York, NY) and pipetted dropwise onto collagen sponges (5 mm diameter, 1 cm height; Kensey Nash/DSM; Exton, PA).

Viability of constructs cultured in αMEM with 10% fetal bovine serum (FBS; Atlanta Biologics; Atlanta, GA) and 1% penicillin-streptomycin-L-glutamine (ThermoFisher) was evaluated using a live/dead assay kit according to manufacturer’s instructions (ThermoFisher). Cylindrical constructs were cut in half lengthwise to expose the center of the sponge, which was imaged.

### Surgical Procedure

Unilateral composite defects were made in 13-week-old female Lewis rats (Charles River Laboratories; Wilmington, MA) by creating an internally stabilized 8 mm defect in the mid-diaphysis of the femur with an overlying 8 mm diameter full-thickness quadriceps defect as previously described ^10^. Collagen sponges loaded with BMP-2 ± MVF were press fit into the defect. Sponges contained 2.5 μg BMP-2 without MVF (n=8 animals), 2.5 μg BMP-2 with MVF (n=8), 10 μg BMP-2 without MVF (n=9), or 10 μg BMP-2 with MVF (n=10). Animal number was determined from previous studies of segmental bone defect healing, and animals were randomly assigned to treatment groups. Animals were double-housed, maintained on a 12-hour light/dark cycle, and allowed ad libitum access to food and water. At 12 weeks post-surgery, animals were euthanized by CO_2_ inhalation. All animal experiments were performed in accordance with the Georgia Institute of Technology Institutional Animal Care and Use Committee (IACUC).

### Bone Regeneration Analyses

Bone healing was qualitatively assessed by radiography at 2, 4, 8, and 12 weeks post-surgery and quantitatively assessed by micro-computed tomography (μCT) at 4, 8, and 12 weeks post-surgery. The defect region was scanned with a voxel size of 38.9 μm. The middle 6.5 mm of the defect was analyzed, and bone volume was quantified by Scanco software using a threshold corresponding to 50% of intact cortical bone ^9; 10^.

Mechanical testing and histology were performed at the 12 week endpoint after euthanasia. Thighs were harvested, soft tissue was cleared, and fixation plates were removed. Femur ends were potted in Wood’s metal (Alfa Aesar) and tested in torsion at a rate of 3°/s to failure (ELF 3200, TA ElectroForce) ^30^. Maximum torque (torque at failure) and torsional stiffness (linear region of torque vs. rotation) were calculated for all samples.

Following mechanical testing, one sample from each group was fixed in 10% neutral buffered formalin (NBF) for 24 hours and decalcified in a formic citrate solution (Newcomer Suppy, Inc; Middleton, WI). Bone and muscle tissues were embedded in paraffin and sectioned at a thickness of 5 μm. Bone tissue was stained with hematoxylin and eosin (H&E); muscle tissue was stained with H&E and with Masson’s Trichrome to observe fibrosis (Histotox Labs; Boulder, CO). Bone sections were also stained with rhodamine-labeled Griffonia simplicifolia lectin I (GS-1 lectin; Vector Laboratories; Burlingame, CA) at a concentration of 10 μg/mL and counterstained with DAPI (ThermoFisher) diluted 1:1000 in phosphate buffered saline to identify blood vessels.

### Serum Cytokine Quantification

At the week 12 endpoint, blood was collected via the rat tail vein from a subset of animals from each treatment group (n=5/group). Serum was collected from whole blood for multiplexed analyte analysis for 27 inflammatory cytokines and chemokines. Samples were prepared using a multiplexed rat cytokine/chemokine magnetic bead panel (MILLIPLEX RECYTMAG-65K), according to manufacturer’s instructions. Data was collected using a MAGPIX multiplexing system (Luminex Corporation; Austin, TX).

### Statistical Analysis

Data were analyzed using GraphPad Prism 7 and are represented as mean ± standard error of the mean (SEM). Bone volume, maximum torque, torsional stiffness, and cytokine levels were analyzed by two-way analysis of variance (ANOVA) and Bonferroni post-hoc tests with an alpha level of 0.05.

## RESULTS

### Construct Viability

MVF cultured in BMP-2 loaded collagen sponges maintained viability in vitro throughout the three-dimensional construct (Figure 1). However, the multicellular MVF appeared to dissociate to single cells by day 3 of culture and reform networks by day 9–14.

**Figure 1.**
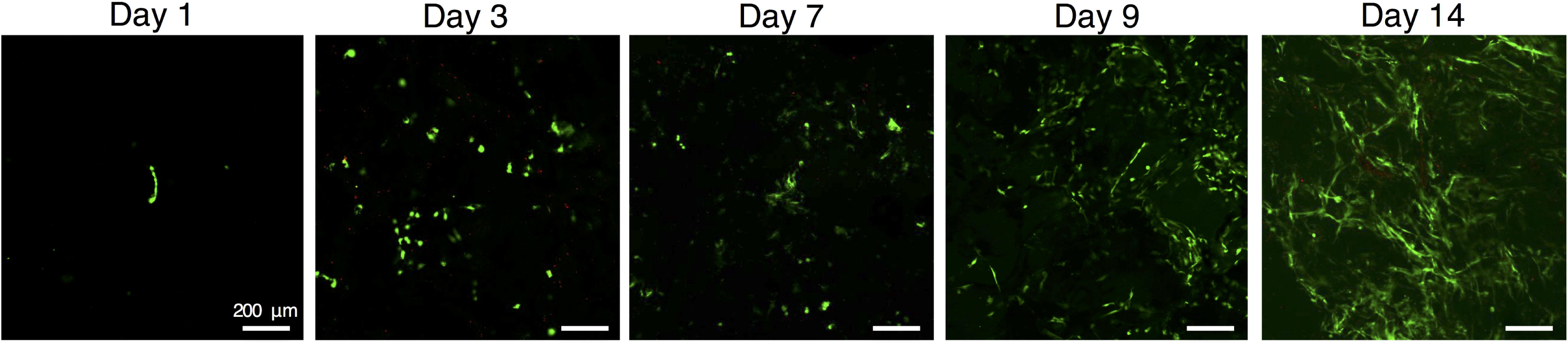
Representative images of MVF cultured in collagen sponge with BMP-2 at days 1, 3, 7, 9, 14. Images were acquired from the center (longitudinal cross section) of the 5 mm diameter sponge. Green, calcein – live, red, ethidium homodimer – dead. MVF dissociated to single cells by day 3 and reformed networks around day 9–14.

### Bone Regeneration

#### Radiography and μCT

Radiographs demonstrated that a majority of defects bridged in all four treatment groups (Figure 2). By 12 weeks, all defects treated with 10 μg BMP-2, with and without MVF, had bridged (nine of nine and ten of ten, respectively). Seven of eight defects treated with 2.5 μg BMP-2 without MVF bridged, and five of eight defects treated with 2.5 μg BMP-2 with MVF bridged. Bone volume in the defect region was measured by μCT and was significantly higher in the 10 μg dose groups than in the 2.5 μg dose groups (Figure 3A; overall effect, p=0.0312). There was no effect of the MVF treatment on bone volume and no significant interaction. The polar moment of inertia was also calculated as a measure of the distribution of bone relative to the central axis. pMOI was significantly lower in the 2.5 μg dose groups (Figure 3B; overall effect, p=0.0047). This corresponds to the more compact morphology observed in the radiographs (Figure 2) and μCT reconstructions of the 2.5 μg dose groups (Figure 3C). Bone density was approximately 870 mm hydroxyapatite/cm^3^ for all four groups.

**Figure 2.**
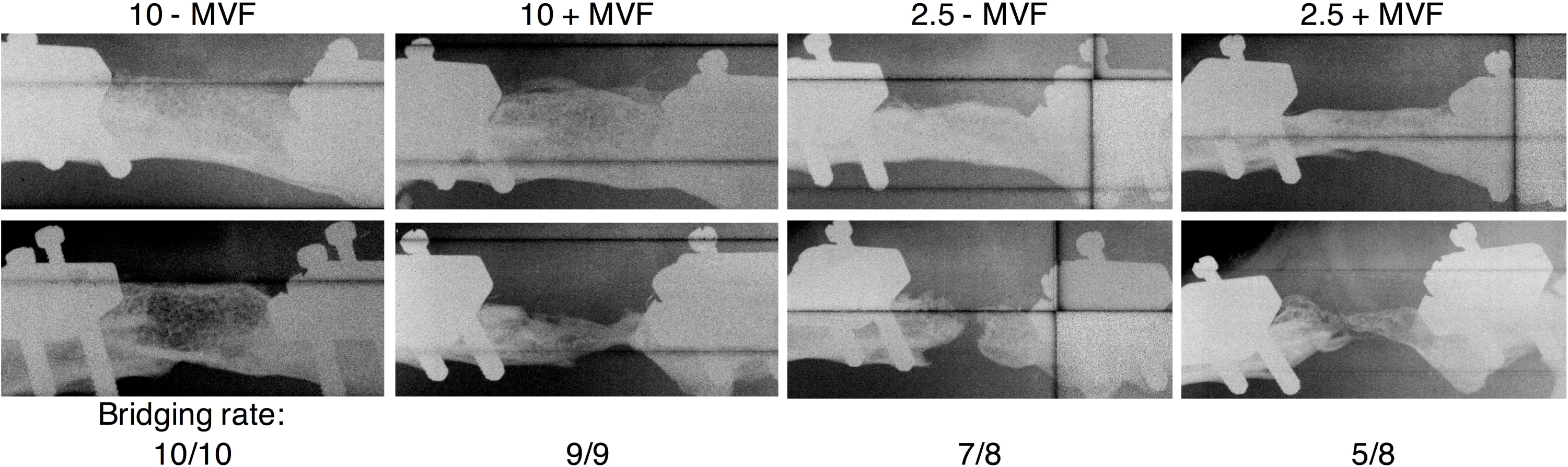
Representative relatively well healed (top) and poorly healed (bottom) 12-week radiographs of each treatment group. Bridging rate by group is shown below.

**Figure 3.**
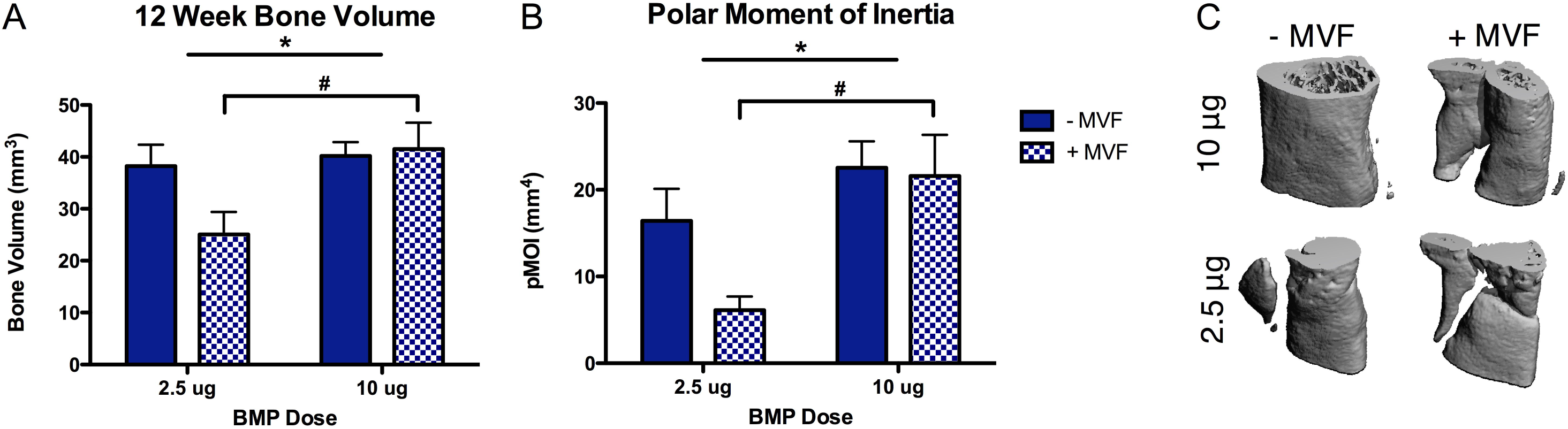
12-week A) bone volume and B) polar moment of inertia of defects as measured by μCT. C) Representative μCT reconstructions of defects from each treatment group that exhibited similar bone volume (~35–40 mm^3^) but different morphologies. * overall effect of BMP dose, p<0.05; # post-hoc p<0.03; 2-way ANOVA; n=8-10/group.

#### Mechanics

The torque to failure was significantly higher for the 10 μg dose BMP-2 groups than the 2.5 μg dose groups (Figure 4; post hoc, p<0.001). There was no effect of MVF and no significant interaction. Despite the increased strength of the 10 μg BMP-2 treatment, the 10 μg dose without MVF group still only achieved 60.5% of intact strength. Stiffness was also significantly higher in the 10 μg dose groups than in the 2.5 μg dose groups (overall effect, p=0.0005), and stiffness was significantly decreased in groups with MVF compared to without MVF (overall effect, p=0.0098).

**Figure 4.**
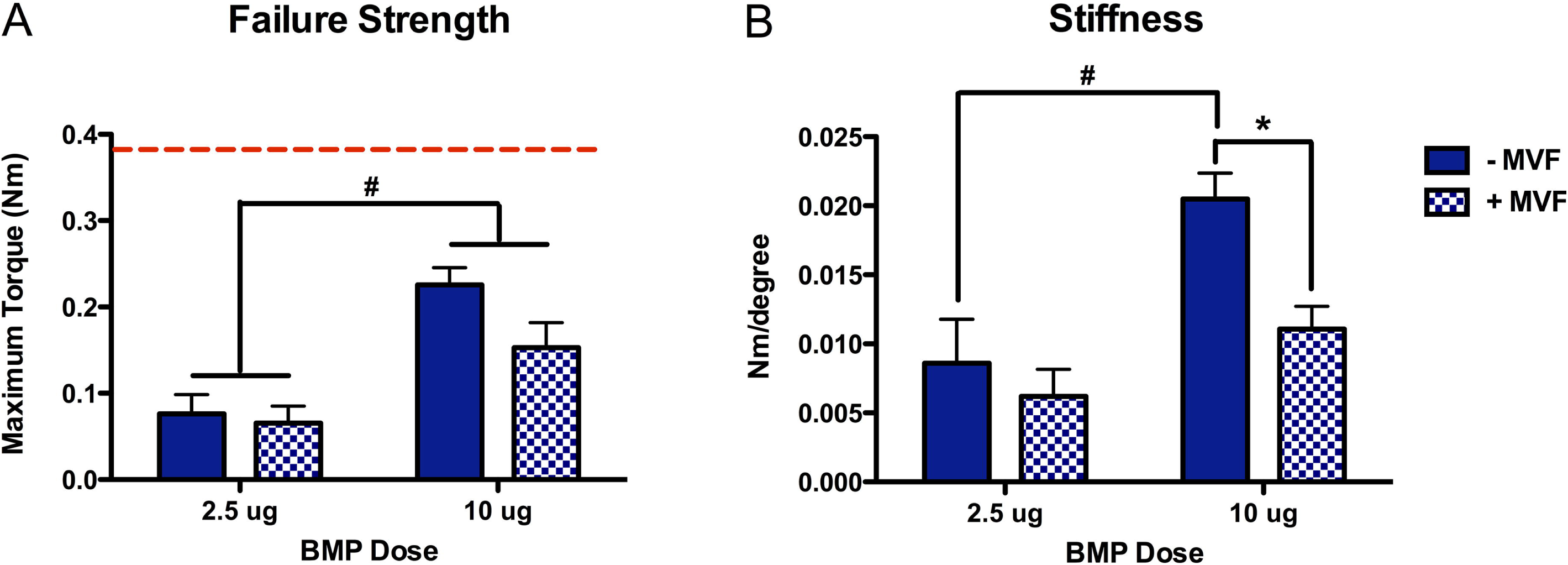
A) Failure strength and B) stiffness of bone regenerated at 12 weeks. Dashed red line indicates failure strength of intact contralateral femur. # post-hoc differences due to BMP dose, p<0.001; * post-hoc differences due to MVF, p<0.01; 2-way ANOVA; n=8-10/group.

### Histology

Qualitative histological assessment of bone defects supported the quantitative μCT data. The 10 μg dose without MVF group showed the greatest amount of bone organized in apparent lamellae, and both groups that received MVF showed larger areas of non-mineralized, marrow-like tissue (Figure 5). There were no qualitative differences in relative number of lectin-stained blood vessel structures among treatment groups at the 12 week time point (Figure 6). As expected, the untreated muscle defects did not heal over the course of this study. At the 12 week end point, muscle exhibited a large degree of fibrosis and fatty infiltration (Figure 7). No qualitative differences in muscle healing were apparent between the well healing 10 μg BMP-2 without MVF and poorly healing 2.5 μg BMP-2 with MVF groups.

**Figure 5.**
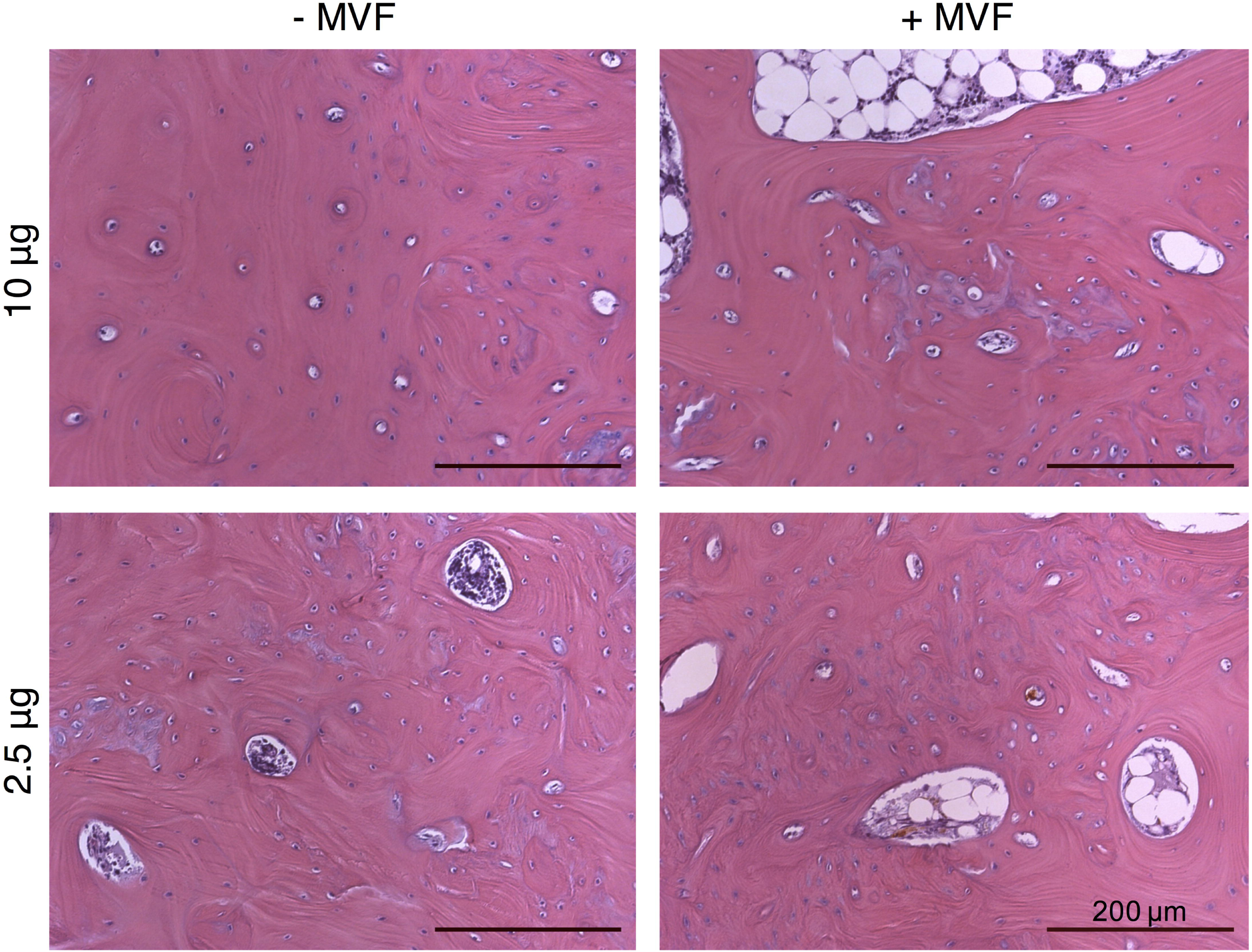
Hematoxylin and eosin staining of bone tissue regenerated by 12 weeks from each treatment group. The 10 μg without MVF group appears the most organized into apparent lamellae, while the other groups are less organized and contain larger areas of non-mineralized, marrow-like tissue.

**Figure 6.**
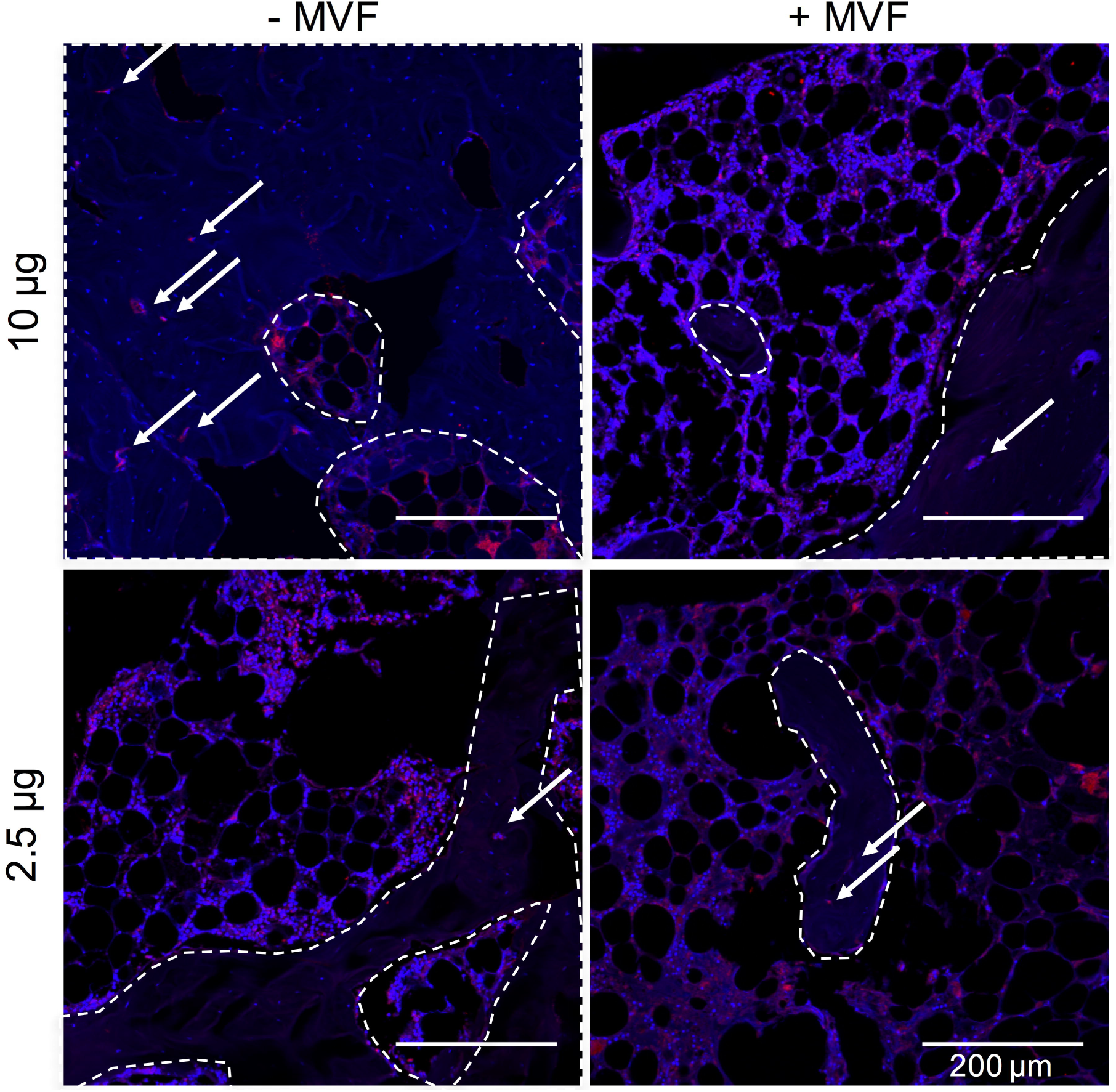
GS-1 lectin staining for blood vessels within regenerate bone at 12 weeks from each treatment group. Dashed line demarcates mineralized vs. marrow-like tissue. White arrows indicate blood vessels within mineralized tissue. Red, GS-1 lectin – vessel glycocalyx; blue, DAPI – nuclei.

**Figure 7.**
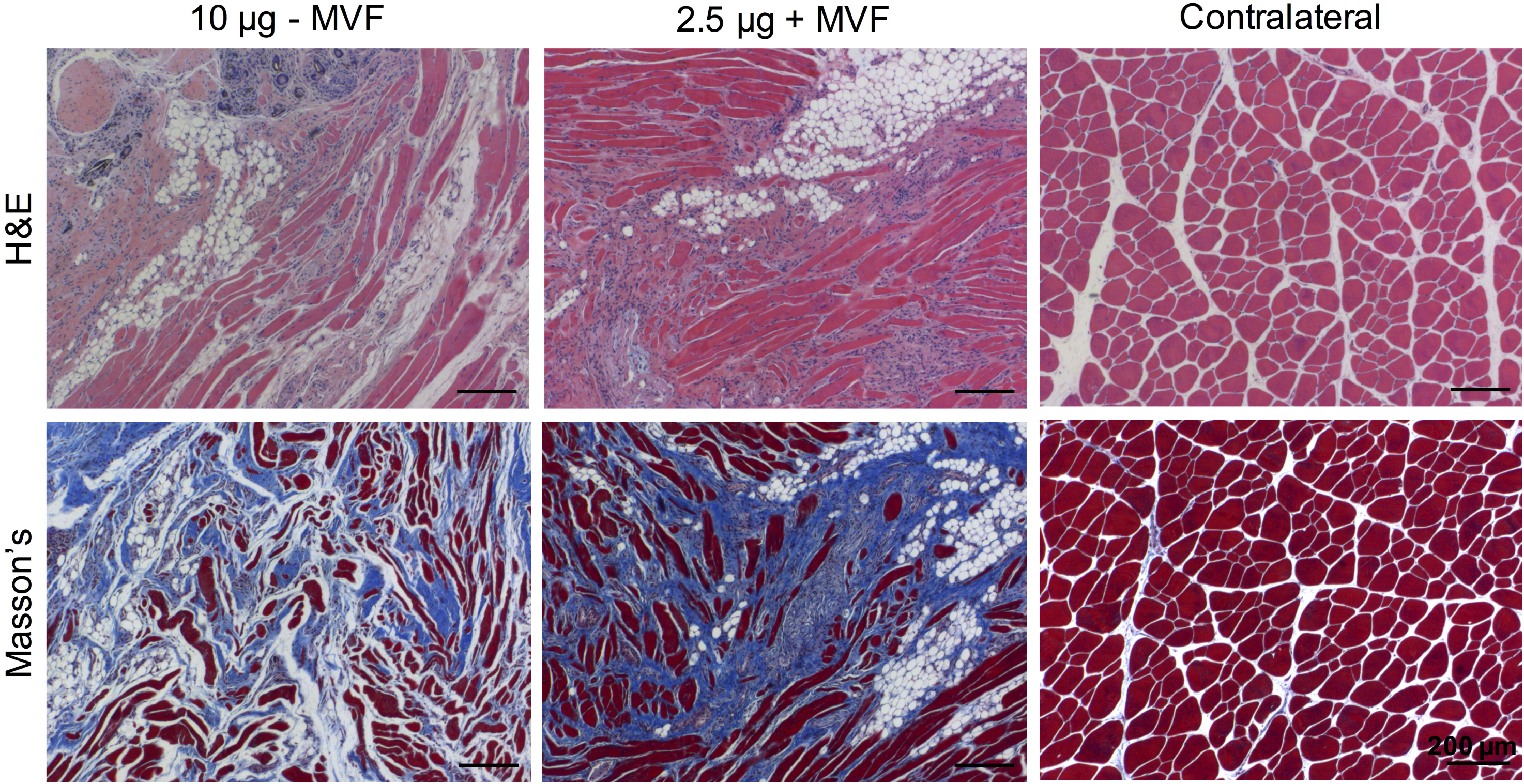
H&E (top) and Masson’s trichrome (bottom; red – muscle fibers, blue – fibrotic tissue) staining of muscle tissue from treatment groups with well healed bone (10 μg - MVF) and poorly healed bone (2.5 μg + MVF) alongside uninjured contralateral muscle tissue.

### Serum Cytokine Quantification

Blood was collected at the 12 week end point to quantify systemic levels of circulating inflammatory cytokines (n=5/group). Levels of multiple interleukins (IL) were significantly decreased in blood samples taken from animals treated with the increased 10 μg BMP-2 dose groups, both with and without MVF (Figure 8). Both pro-inflammatory interleukins, such as IL-1a and IL-1b, and anti-inflammatory interleukins, such as IL-5, IL-10, and IL-13, ^31^ were decreased in the 10 μg BMP-2 dose groups Pro-inflammatory eotaxin was also lower (p=0.0507) in the 10 μg BMP dose groups.

**Figure 8.**
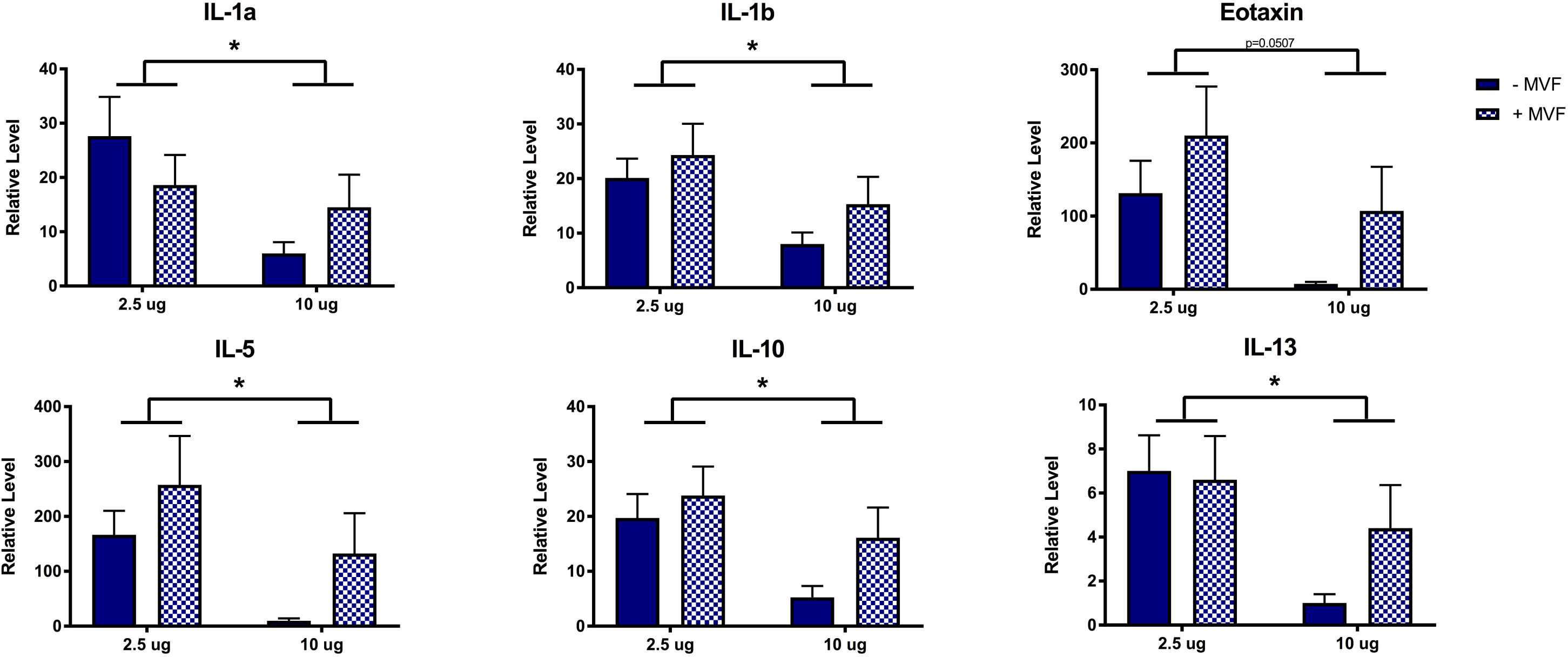
Relative serum levels of pro- (top) and anti-inflammatory (bottom) cytokines by treatment group as measured by multiplexed analyte analysis. * post-hoc differences due to BMP dose, p<0.05; 2-way ANOVA; n=5/group.

## DISCUSSION

Composite tissue extremity injuries represent a significant challenge to orthopaedic surgeons and a momentous hardship for patients. Vascularization is an enabling step of wound healing, and we hypothesized that BMP-2-mediated functional regeneration of composite extremity injuries would be dose dependent and could be further enhanced via co-delivery of adipose-derived microvascular fragments (MVF), which have been previously shown to increase tissue vascular volume.

BMP-2 is clinically approved for a number of bone healing indications and has shown success in treating long bone fractures and non-unions ^6; 32^. There is a BMP-2 dose response in humans and in pre-clinical animal models, but precise dosing required for specific human applications is not well defined ^9; 32^. High BMP-2 dosage has been linked a number of side effects including heterotopic ossification (HO) and increased inflammation ^7^, and HO is also often increased in cases of trauma ^33–35^. While a 2.5 μg dose of BMP-2 is a bridging dose in a segmental defect model ^9^, it produces inconsistent bridging in this challenging composite defect model ^10^. In this segmental defect model, HO has been observed at 30 μg doses of BMP-2 ^36; 37^. We showed in the present study that a 10 μg dose is sufficient to improve composite defect healing relative to the 2.5 μg dose and does not lead to mineralization outside the collagen sponge delivery vehicle. A moderately increased dose (e.g. 10 μg relative to 2.5 μg or 30 μg) may be required in composite injuries to compensate for the loss of endogenous stem and progenitor cells and growth factor availability as a result of the concomitant muscle loss. However, these studies together demonstrate the need for precise identification of the appropriate dosing window of BMP-2. While muscle loss constitutes an increased level of trauma, this model does not incorporate head trauma or muscle crush, which are known contributors to human post traumatic HO ^34; 35^. Traumatically injured muscle can give rise to osteogenic progenitor cells ^35^, but it is unknown whether a similar effect would be present in this model which cleanly removes a volumetric section of muscle with a biopsy punch. While large volume muscle loss has been shown to have a more profound impact on bone healing than muscle crush injuries ^38^, the VML model utilized here may not fully recapitulate the HO risk associated with trauma of clinical open fractures that often also have significant muscle damage. Future work may be warranted to investigate more aggressive debridement of damaged muscle adjacent to bone injuries as prophylaxis for HO, though this could potentially lead to reduced functional outcomes.

Local healing at the site of a regenerative intervention, as well as HO, are both thought to be influenced by the systemic immune response. Interestingly, the 10 μg BMP dose resulted in lower levels of both pro- and anti-inflammatory systemic cytokines relative to the 2.5 μg dose at the 12-week time point. The 2.5 μg dose group’s failure to completely heal may have prolonged the inflammatory stage of the wound healing process, which previous studies have shown to be increased in composite tissue injuries ^27^. That persistent local inflammation may then have resulted in altered systemic immune cytokine profiles. Rather than a high BMP dose inducing inflammation, the successful bridging and restoration of function may have resolved the local inflammation and led to lower cytokine levels by the 12 week time point. Future studies investigating inflammatory profiles at earlier time points would be required to more definitively address the composite injury’s inflammatory response to various BMP doses.

MVF was used as a proposed therapeutic approach to enhance revascularization and subsequently bone regeneration; however, the MVF treatment did not improve bone volume or failure strength and in fact decreased the stiffness of the regenerated bone. Although MVF treatment had no significant effect on bone volume, the mineralization pattern of the regenerated bone appeared more fragmented in MVF-treated defects as observed with μCT. The fragmented mineralization of MVF-treated groups may have contributed to the differences in mechanical properties.

Delivery of MVF did not result in qualitative differences in lectin-stained blood vessel structures at 12 weeks. Cell implantation strategies broadly are challenged by issues of viability and retention ^39–41^. Although vascularization is hypothesized to improve viability by enabling nutrient and oxygen transport, the implanted MVF may have failed to survive long enough to inosculate with the host vasculature. In this study, while in vitro viability of MVF was maintained in collagen sponge, the multicellular MVF first dissociated to single cells and then reformed networks. This is not consistent with the sprouting angiogenesis typically observed in MVF within a collagen gel ^20; 24; 42–44^ and may have contributed to the lack of effect of MVF on bone healing. While collagen sponge is a clinically available BMP-2 delivery vehicle, it may not be an optimal delivery vehicle for MVF, which are exquisitely sensitive to extracellular matrix properties ^42; 45^. Further, MVF exert traction forces on their matrices and can contract collagen hydrogels ^20; 44; 46^; cell-mediated contraction of collagen sponge may have contributed to the more fragmented mineralization observed in the MVF-treated defects (Fig 3C). Future studies to assess MVF retention, viability, and vascularization at earlier time points are required.

Beyond the vascular components of adipose-derived MVF, these structures also contain mesenchymal stem cells (MSCs) ^26; 47^. Exogenously delivered adipose-derived MSCs have previously been shown to negatively affect mineralization when co-delivered with BMP-2 ^48^. While BMP-2 upregulates osteogenic signaling of bone marrow-derived MSCs, adipose-derived MSCs may differentially interact with BMP-2 to dampen or alter its signaling to endogenous host cells. However, adipose-derived MVF contain only about 2%-7% MSCs ^26; 49^, so their relative contribution may be proportionally small. Retention of other adipose-derived cells may have contributed to the larger marrow spaces histologically observed in MVF-treated defects (Figure 5).

This is the first investigation of BMP-2 dose in a composite defect model. While the delivery of MVF did not improve bone healing in the composite defect model, an increased dose of 10 μg BMP-2 significantly improved bone healing and regenerate bone mechanics relative to a 2.5 μg dose. While 2.5 μg BMP-2 restored bone failure strength to only 20% of intact strength, a 10 μg BMP dose restored failure strength to 60% of intact. This improvement may represent significant functional improvements but also underscores the challenging nature of healing composite tissue injuries. Collagen sponge is a clinically approved, and therefore translationally relevant, BMP-2 delivery vehicle but may not be an appropriate MVF delivery vehicle. A modest increase in BMP-2 dosage may be a clinically relevant treatment strategy for critical size bone defects with concomitant muscle damage while a biomaterial delivery system for co-delivery of MVF is developed. Approaches that enhance early vascularization may be required to restore regenerated bone to greater than 60% of intact strength in cases of composite bone and muscle traumatic injury.

## ACKNOWLEDGEMENTS

This work was supported by funding from the Armed Forces Institute for Regenerative Medicine (AFIRM II) effort under award number W81XWH-14-2-0003. Opinions, interpretations, conclusions, and recommendations are those of the authors and are not necessarily endorsed by the Department of Defense. This work also received funding from the Center for Regenerative Engineering and Medicine (REM) research center seed grant program, supported by the National Center for Advancing Translational Sciences of the National Institutes of Health under Award Number UL1TR00454. The content is solely the responsibility of the authors and does not necessarily represent the official views of the National Institutes of Health. The authors wish to acknowledge the core facilities at the Parker H. Petit Institute for Bioengineering and Bioscience at the Georgia Institute of Technology for the use of their shared equipment, services, and expertise. We also thank Ryan Akman, Olivia Burnsed, Albert Cheng, Gilad Doron, Brett Klosterhoff, Lina Mancipe-Castro, David Reece, Giuliana Salazar-Noratto, and Brennan Torstrick for their assistance with surgeries and Levi Wood for usage of the MAGPIX multiplexing system.

Author contributions
MAR, LK, CEV, and YW performed experiments. MAR, LK, HYS, REG, and NJW contributed to data interpretation. MAR, LK, KR, REG, and NJW designed experiments. All authors have read and approved the final submitted manuscript.

## REFERENCES

1. Zura R, Xiong Z, Einhorn T, et al. 2016. Epidemiology of Fracture Nonunion in 18 Human Bones. JAMA Surg 151:e162775.

2. Hak DJ, Fitzpatrick D, Bishop JA, et al. 2014. Delayed union and nonunions: epidemiology, clinical issues, and financial aspects. Injury 45 Suppl 2:S3–7.

3. MacKenzie EJ, Bosse MJ, Pollak AN, et al. 2005. Long-term persistence of disability following severe lower-limb trauma. Results of a seven-year follow-up. J Bone Joint Surg Am 87:1801–1809.

4. Boden SD, Kang J, Sandhu H, et al. 2002. Use of recombinant human bone morphogenetic protein-2 to achieve posterolateral lumbar spine fusion in humans: a prospective, randomized clinical pilot trial: 2002 Volvo Award in clinical studies. Spine (Phila Pa 1976) 27:2662–2673.

5. Carragee EJ, Hurwitz EL, Weiner BK. 2011. A critical review of recombinant human bone morphogenetic protein-2 trials in spinal surgery: emerging safety concerns and lessons learned. Spine J 11:471–491.

6. Govender S, Csimma C, Genant HK, et al. 2002. Recombinant human bone morphogenetic protein-2 for treatment of open tibial fractures: a prospective, controlled, randomized study of four hundred and fifty patients. J Bone Joint Surg Am 84-A:2123–2134.

7. James AW, LaChaud G, Shen J, et al. 2016. A Review of the Clinical Side Effects of Bone Morphogenetic Protein-2. Tissue engineering Part B, Reviews 22:284–297.

8. Nauth A, Giles E, Potter BK, et al. 2012. Heterotopic ossification in orthopaedic trauma. Journal of orthopaedic trauma 26:684–688.

9. Boerckel JD, Kolambkar YM, Dupont KM, et al. 2011. Effects of protein dose and delivery system on BMP-mediated bone regeneration. Biomaterials 32:5241–5251.

10. Willett NJ, Li MT, Uhrig BA, et al. 2013. Attenuated human bone morphogenetic protein-2-mediated bone regeneration in a rat model of composite bone and muscle injury. Tissue engineering Part C, Methods 19:316–325.

11. Gustilo RB, Merkow RL, Templeman D. 1990. The management of open fractures. J Bone Joint Surg Am 72:299–304.

12. Einhorn TA, Gerstenfeld LC. 2015. Fracture healing: mechanisms and interventions. Nat Rev Rheumatol 11:45–54.

13. Hyzy SL, Kajan I, Wilson DS, et al. 2017. Inhibition of angiogenesis impairs bone healing in an in vivo murine rapid resynostosis model. Journal of biomedical materials research Part A 105:2742–2749.

14. Lu C, Miclau T, Hu D, et al. 2007. Ischemia leads to delayed union during fracture healing: a mouse model. Journal of orthopaedic research : official publication of the Orthopaedic Research Society 25:51–61.

15. Garcia JR, Clark AY, Garcia AJ. 2016. Integrin-specific hydrogels functionalized with VEGF for vascularization and bone regeneration of critical-size bone defects. Journal of biomedical materials research Part A 104:889–900.

16. Patel ZS, Young S, Tabata Y, et al. 2008. Dual delivery of an angiogenic and an osteogenic growth factor for bone regeneration in a critical size defect model. Bone 43:931–940.

17. Subbiah R, Hwang MP, Van SY, et al. 2015. Osteogenic/angiogenic dual growth factor delivery microcapsules for regeneration of vascularized bone tissue. Adv Healthc Mater 4:1982–1992.

18. Wang L, Fan H, Zhang ZY, et al. 2010. Osteogenesis and angiogenesis of tissue-engineered bone constructed by prevascularized beta-tricalcium phosphate scaffold and mesenchymal stem cells. Biomaterials 31:9452–9461.

19. Zhang W, Zhu C, Wu Y, et al. 2014. VEGF and BMP-2 promote bone regeneration by facilitating bone marrow stem cell homing and differentiation. Eur Cell Mater 27:1–11; discussion 11–12.

20. Hoying JB, Boswell CA, Williams SK. 1996. Angiogenic potential of microvessel fragments established in three-dimensional collagen gels. In vitro cellular & developmental biology Animal 32:409–419.

21. Laschke MW, Menger MD. 2015. Adipose tissue-derived microvascular fragments: natural vascularization units for regenerative medicine. Trends in biotechnology 33:442–448.

22. Chang CC, Krishnan L, Nunes SS, et al. 2012. Determinants of microvascular network topologies in implanted neovasculatures. Arteriosclerosis, thrombosis, and vascular biology 32:5–14.

23. Nunes SS, Greer KA, Stiening CM, et al. 2010. Implanted microvessels progress through distinct neovascularization phenotypes. Microvascular research 79:10–20.

24. Li MT, Ruehle M, Stevens H, et al. 2017. Skeletal myoblast-seeded vascularized tissue scaffolds in the treatment of a large volumetric muscle defect in the rat biceps femoris muscle. Tissue engineering Part A.

25. Frueh FS, Spater T, Korbel C, et al. 2018. Prevascularization of dermal substitutes with adipose tissue-derived microvascular fragments enhances early skin grafting. Scientific reports 8:10977.

26. Frueh FS, Spater T, Lindenblatt N, et al. 2017. Adipose Tissue-Derived Microvascular Fragments Improve Vascularization, Lymphangiogenesis, and Integration of Dermal Skin Substitutes. J Invest Dermatol 137:217–227.

27. Hurtgen BJ, Ward CL, Garg K, et al. 2016. Severe muscle trauma triggers heightened and prolonged local musculoskeletal inflammation and impairs adjacent tibia fracture healing. J Musculoskelet Neuronal Interact 16:122–134.

28. Pollot BE, Goldman SM, Wenke JC, et al. 2016. Decellularized extracellular matrix repair of volumetric muscle loss injury impairs adjacent bone healing in a rat model of complex musculoskeletal trauma. J Trauma Acute Care Surg 81:S184–S190.

29. Hurtgen BJ, Henderson BEP, Ward CL, et al. 2017. Impairment of early fracture healing by skeletal muscle trauma is restored by FK506. BMC musculoskeletal disorders 18:253.

30. Oest ME, Dupont KM, Kong HJ, et al. 2007. Quantitative assessment of scaffold and growth factor-mediated repair of critically sized bone defects. Journal of orthopaedic research : official publication of the Orthopaedic Research Society 25:941–950.

31. Opal SM, DePalo VA. 2000. Anti-inflammatory cytokines. Chest 117:1162–1172.

32. Gautschi OP, Frey SP, Zellweger R. 2007. Bone morphogenetic proteins in clinical applications. ANZ J Surg 77:626–631.

33. Potter BK, Burns TC, Lacap AP, et al. 2007. Heterotopic ossification following traumatic and combat-related amputations. Prevalence, risk factors, and preliminary results of excision. J Bone Joint Surg Am 89:476–486.

34. Cipriano CA, Pill SG, Keenan MA. 2009. Heterotopic ossification following traumatic brain injury and spinal cord injury. J Am Acad Orthop Surg 17:689–697.

35. Jackson WM, Aragon AB, Bulken-Hoover JD, et al. 2009. Putative heterotopic ossification progenitor cells derived from traumatized muscle. Journal of orthopaedic research: official publication of the Orthopaedic Research Society 27:1645–1651.

36. Krishnan L, Priddy LB, Esancy C, et al. 2017. Delivery vehicle effects on bone regeneration and heterotopic ossification induced by high dose BMP-2. Acta biomaterialia 49:101–112.

37. Hettiaratchi MH, Chou C, Servies N, et al. 2017. Competitive Protein Binding Influences Heparin-Based Modulation of Spatial Growth Factor Delivery for Bone Regeneration. Tissue engineering Part A 23:683–695.

38. Utvag SE, Grundnes O, Rindal DB, et al. 2003. Influence of extensive muscle injury on fracture healing in rat tibia. Journal of orthopaedic trauma 17:430–435.

39. Allen AB, Zimmermann JA, Burnsed OA, et al. 2016. Environmental manipulation to promote stem cell survival in vivo: use of aggregation, oxygen carrier, and BMP-2 co-delivery strategies. J Mater Chem B 4:3594–3607.

40. Ruehle MA, Stevens HY, Beedle AM, et al. 2018. Aggregate mesenchymal stem cell delivery ameliorates the regenerative niche for muscle repair. Journal of tissue engineering and regenerative medicine.

41. Levit RD, Landazuri N, Phelps EA, et al. 2013. Cellular encapsulation enhances cardiac repair. Journal of the American Heart Association 2:e000367.

42. Krishnan L, Hoying JB, Nguyen H, et al. 2007. Interaction of angiogenic microvessels with the extracellular matrix. American journal of physiology Heart and circulatory physiology 293:H3650–3658.

43. Krishnan L, Underwood CJ, Maas S, et al. 2008. Effect of mechanical boundary conditions on orientation of angiogenic microvessels. Cardiovasc Res 78:324–332.

44. Ruehle MA, Krishnan L, LaBelle SA, et al. 2017. Decorin-containing collagen hydrogels as dimensionally stable scaffolds to study the effects of compressive mechanical loading on angiogenesis. MRS Communications 7:466–471.

45. Edgar LT, Hoying JB, Utzinger U, et al. 2014. Mechanical interaction of angiogenic microvessels with the extracellular matrix. Journal of biomechanical engineering 136:021001.

46. Edgar LT, Underwood CJ, Guilkey JE, et al. 2014. Extracellular matrix density regulates the rate of neovessel growth and branching in sprouting angiogenesis. PloS one 9:e85178.

47. Laschke MW, Grasser C, Kleer S, et al. 2014. Adipose tissue-derived microvascular fragments from aged donors exhibit an impaired vascularisation capacity. Eur Cell Mater 28:287–298.

48. Dosier CR, Uhrig BA, Willett NJ, et al. 2015. Effect of cell origin and timing of delivery for stem cell-based bone tissue engineering using biologically functionalized hydrogels. Tissue engineering Part A 21:156–165.

49. Laschke MW, Karschnia P, Scheuer C, et al. 2017. Effects of cryopreservation on adipose tissue-derived microvascular fragments. Journal of tissue engineering and regenerative medicine.

